# Shifting sensitivity and signal-dependent timing in the copulatory displays of songbirds

**DOI:** 10.1101/2021.05.19.444794

**Authors:** Ammon Perkes, Bernd Pfrommer, Kostas Daniilidis, David White, Marc Schmidt

**Affiliations:** University of Pennsylvania, Philadelphia, Pennsylvania, United States; University of California Davis, Davis, California, United States; Wilfrid Laurier University, Waterloo, Ontario, Canada

## Abstract

Female preferences for male signals largely determine courtship success. In most songbirds, females control reproduction via the copulation solicitation display (CSD), an innate, stereotyped posture produced in direct response to male displays. In some species, CSD can be elicited under controlled experimental conditions, allowing CSD production to be used as a bioassay for the underlying mechanics of courtship, however historically our ability to measure variability in CSD was limited. Using computer vision approaches to precisely quantify the trajectory of CSD in female brown-headed cowbirds (*Molothrus ater*), we show that song potency (how often a song elicits CSD) and female responsivity (how likely a female is to produce CSD) each predict whether a female produces CSD in response to a given presentation. Within elicited CSD, potency and responsivity also predict the duration and onset latency of the displays. By modeling and testing two alternative hypotheses of preference formation, we show that changes in response rates are consistent with a shifting receiver response threshold, suggesting that the song quality required to evoke the copulatory posture in a particular female will vary over time. These results establish a novel framework for exploring CSD and emphasize the importance of the internal state of receivers in determining behavioral responses and suggest that female responsivity acts in conjunction with male signal strength to determine the efficacy of male courtship signaling.

## Introduction

Courtship signals are usually, by necessity, easily observed. Receiver preference, in contrast, is hidden, except as manifested by some detectable behavioral response. Because of this difficulty in precisely quantifying reception, our understanding of receivers is outpaced by our understanding of signalers for both neurobiology and evolutionary function. While researchers have made meaningful progress in identifying male traits that predict their courtship success (i.e., being “chosen” by a female) (Andersson 1994; Reaney and Backwell 2007; Keagy et al. 2011), we know less about the causal relationships between these traits, female preference, and fitness (Reid et al. 2004; Faust and Goldstein 2021; Godin and Dugatkin 1996), and “embarrassingly little” about the female mechanisms that mediate mate choice (Deangelis and Hofmann 2020). These gaps severally limit our understanding of courtship in general and the action of female preference in particular, leaving us with only one half of the equation of reproductive success. Fully understanding courtship requires developing precise behavioral assays and analytical tools to investigate the mechanisms of female preference and response, similar to the well used acoustic techniques for quantifying male signals.

While it is possible to monitor females in natural contexts and observe with whom they mate (Keagy et al. 2011; Faust and Goldstein 2021), experimental approaches require methods that can control for the myriad factors that influence female choice. As such, most of our understanding of courtship comes from laboratory tests studying proxies for female preference, such as place-preference tasks (Riebel and Slater 1998; R. C. Anderson 2009; MacLaren and Rowland 2006; Barr et al. 2021), phonotaxis (Ryan 1985; Morris et al. 1978) and others (King and West 1977; Rodriguez-Sierra et al. 1975; Hardy and DeBold 1971). Songbirds provide an effective system for studying female preferences because female mate selection is largely based on preferences for male song, and females produce an overt, visual response corresponding to their preference, the Copulation Solicitation Display (CSD). In many songbird species, copulation requires the female to produce this lordosis-like pose, such that it is not only a signal of preference but a key behavioral gate on mating. During this display, females spread their wings, raise their tail feathers, and arch their backs to expose their cloaca (Video S1). Female CSD invites, facilitates, and controls access to copulation (King and West 1977; Searcy and Marler 1981; Catchpole et al. 1986), such that, for males, eliciting a CSD is critical for reproductive success. While in some species, CSD is only observed in natural contexts, in many species it is possible to elicit CSD using only playback of male song, allowing the stimulus and response to be studied under controlled conditions (King and West 1977; Searcy and Marler 1981; Kreutzer et al. 1992). Furthermore, in these contexts, how often females produce CSD to a given song serves as a natural bioassay of the signal strength, since some songs are more potent in eliciting CSD. These potency scores, measured in CSD assays, correspond to the preferences measured in natural contexts (O’Loghlen and Rothstein 1995; West et al. 1981), suggesting they reflect female preferences and the efficacy of song as they occur in the wild.

Studies using the CSD essay have been extraordinarily effective at investigating mechanisms of female preferences and the features male song (King and West 1977; Searcy and Marler 1981; Kreutzer et al. 1992; Brenowitz 1991; Maguire et al. 2013). Unfortunately, often it was only possible to document whether or not a female produced a CSD. Additionally, because these experiments were space and time intensive, dataset size necessarily limited and this approach fails to capture any level of variation in the display. Rarely, researchers have manually observed videos to provide descriptions of CSD intensity or duration (O’Loghlen and Rothstein 1995, 2010a), but these measures still pale in comparison to the temporal and spatial precision with which we can measure elements of song, leaving us lacking in knowledge regarding the fine scale tuning between song characteristics and female preferences, individual differences in responsivity, neural control, plasticity and learning effects, and other aspects of female preferences. The goal of this work is to develop a new system to study the CSD at a high level of precision. Here, using brown-headed cowbirds, we develop a computer vision tracking system that can automatically quantify variation within the timing and intensity of CSD at a level of precision and through-put previously impossible, enabling both fine-scale quantification of responses during playback and longer-term comparisons of the changes to behavior as females hear songs multiple times and potentially change their assessments of songs with experience.

First, we explore the temporal and postural dynamics of CSD reactivity, examining consistency and variation both within and across females. Just as song has a consistent structure with meaningful variation within and among songs (Catchpole and Slater 2003), there could be variation in the timing, intensity, and the specific components of CSD, similar to other postural behaviors in courtship (Hebets and Uetz 2000; Fusani et al. 1997; Mitoyen et al. 2019; O’Loghlen and Rothstein 2010b) that are critical for mating success. We quantify this variation and test whether variation in the timing and extent of CSD is linked to the potency of song and the responsivity of females, providing insight in the possible mechanisms that underly CSD production.

Second, we assess how female CSD response changes over the course of many trials. During the breeding season in nature, females hear up to thousands of songs over the course of the day, but CSDs occur much more rarely. We know very little about what determines when females produce CSD, but existing research (both in cowbirds and other species) clearly demonstrates that a) CSD is gated by the hormonal state of the female, such that CSD only occur naturally during breeding season (White et al. 2006) or based on exogenous estradiol implants (O’Loghlen and Rothstein 2012), and b) that female preferences are plastic and change with exposure to song (Riebel 2003; Miller 1979; Lauay et al. 2004; Vyas et al. 2009; Wall and Woolley 2024), tutoring (Freed-Brown and White 2009) and the other song sets within a playback session (West and King 1988). It seems likely then that these two variable mechanisms—female receptivity and song preference—combine to determine whether CSD is produced. To test how these mechanisms interact, we first develop a signal-response model in order to generate explicit predictions. Then we take advantage of a novel approach for generating high-throughput song playbacks, paired with many years of existing playback data, to assess if and how song preferences and song responsivity vary over playbacks. By describing variation within CSD and the identifying the underlying features which predict it, we gain insight into the degree of control females have during CSD production and the mechanisms which likely drive this behavior.

## Results

We combined data from several prior playback experiments with additional experiments performed between 2018-2019 (Figure 1) to build a large dataset of song responses. Within the dataset, baseline response rates varied (Figure S1), but both within and across experimental groups, females showed clear and consistent preference for some songs over others (Permutation test, p < 0.0001). For example, in the most responsive group, the two most successful songs elicited CSD for over 75% of song-playbacks while the remaining 8 songs (i.e., those less likely to evoke CSD) elicited CSD in less than 41% of playbacks total (based on n=47 presentations per song, spread across 6 birds). Potency rankings were consistent within experimental groups (mean pair-wise Pearson’s r = 0.193 +- 0.02, SE), as well as across experimental groups of birds for which we used the same song set (for group averages, mean Pearson’s r = 0.414, +- 0.06, SE). While preferences were broadly consistent, we observed clear differences in receiver responsiveness (Anova, F = 101.7, p < .0001, Figure S1). Each female’s response rate was calculated at the time of playback and this varied drastically within and across birds. For example, the most responsive female produced CSD in response to 100% of song playbacks (n=104), while 22 out of 68 females responded to no more than 10% of song presentations. Responsiveness also varied within birds, with individuals often producing CSD in response to 100% of songs in some playback blocks but 0% of songs in others.

**Figure 1.**
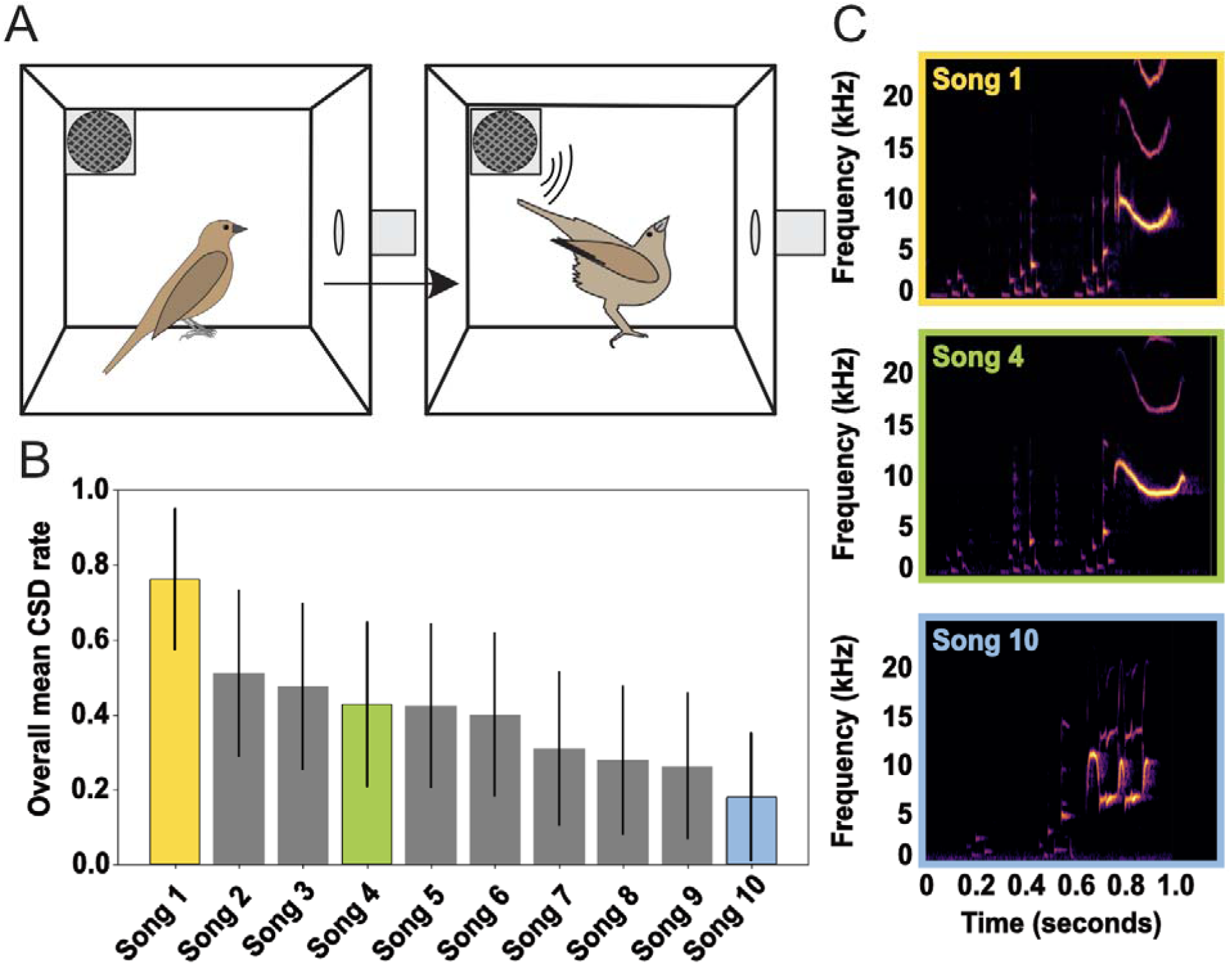
Experimental paradigm for evaluating CSD responsiveness. A. Isolated adult female cowbirds in breeding condition placed in a sound isolation chamber produce CSDs in response to playback of recorded male song. B. Song potency ranking based on the presentation of a battery of 10 songs, with example songs in C. Each song was presented 10 times in randomized blocks, with presentations separated by 90 minutes, over a period of 2 weeks) to n=5 females. Comparison of mean response rate across all presentations provides the relative song potency of each song.

These two measures, song potency and female responsivity, are central to our subsequent results, which we divide according to our two specific questions: In <u>part 1</u>, we use image analysis and computer vision approaches to quantify female pose and variation across CSDs, testing whether variation in CSD is linked to song potency and/or female responsivity. In <u>part 2</u>, we test whether song preference formation in our CSD assay is better explained by a weight-shift model (i.e., shifting potency) or a threshold-shift model (i.e., shifting responsivity) by using agent-based simulations to identify clear predictions that can reliably distinguish between the two hypotheses, and then testing our data against these predictions.

### Part 1: The timing and trajectory of CSD varies within and across individual and song

By definition, song potency describes how likely a song is to elicit a CSD, while responsivity estimates how likely a female is to produce a CSD; these metrics—as proxies for song quality and female state respectively—may also predict the timing and trajectory of song, such that the “best” songs and/or females also produce lower latency, higher duration, and/or more dramatic CSDs. Using multiple cameras in a calibrated enclosure, we were able to precisely reconstruct 3d bird pose during CSD production (Video S2) and measure latency, duration, and behavioral trajectory. Here we test how these metrics varied across CSD responses as a function of song potency and estimated receiver responsiveness.

### Song potency and receiver responsiveness both predict onset latency

Within our multi-camera enclosures, we calculated onset latency for 8 birds (Figure 2A). Mean latency for the 268 CSDs we measured was 0.628 s (± 0.249), shorter than the duration of song, with significant individual variation between birds (Anova, F=88.34 p<.0001). Using linear mixed models to account for song and bird identity, we found a significant effect of song potency on CSD latency (Figure 2B, e=-0.230, CI: −0.373:-0.091, p<.005) indicating that (after controlling for individual responsivity) the highest potency songs elicited responses 230 ms earlier than the lowest potency songs. As with song potency, responsivity, i.e., the mean responsiveness within a given presentation block, had a significant effect on latency (Figure 2C, e=-0.113, CI: −0.213:-0.014, p<.05) with CSD response latencies being faster in trials when females were more responsive (i.e., had a lower response threshold). This effect corresponds to a 113 ms difference in latency for high-responsive vs low-responsive females.

**Figure 2.**
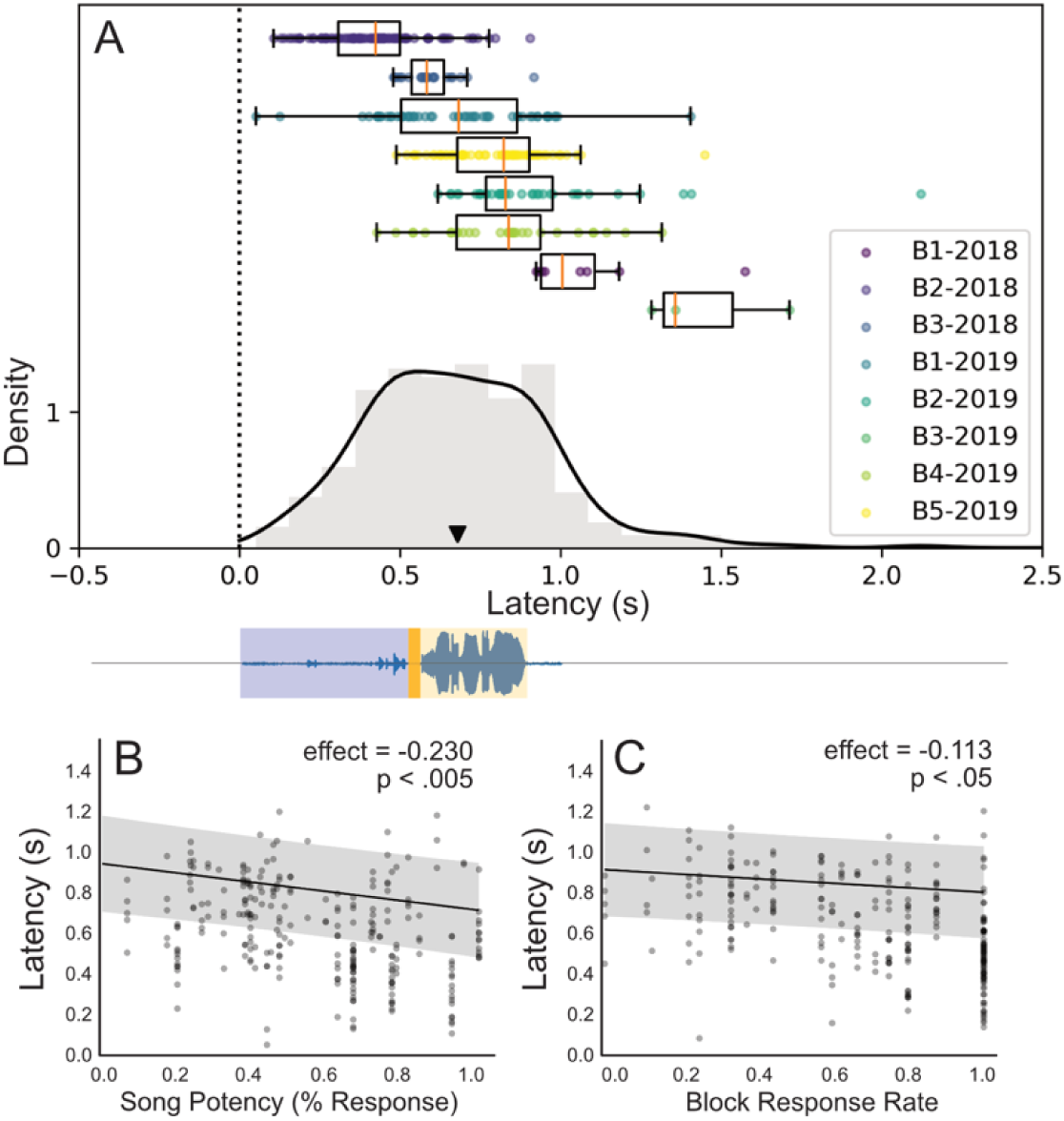
Latency varies with both song potency and underlying receiver state. **A.** The distribution of CSD latencies for 268 CSDs from 8 individual birds, sorted by latency and aligned to song onset. The blue and orange shading shows the song with the transition between colors highlighting the transition from introductory notes to terminal whistle. Note that introductory notes are much lower amplitude, compared to the whistle. **B.** Using a linear mixed model, we show that latency to CSD onset decreases with song potency, with a 230 ms difference in the latency between the highest and lowest potency songs. Each point represents one CSD and the black line shows the linear model estimate. C. Latency to CSD onset also decreases as a function of the female’s block response rate, i.e., the proportion of presentations which elicit a CSD in one set of (usually ten) song presentations, with highly responsive females producing CSDs 113ms sooner than low-responsive females. As in B, each point represents a single CSD, and the black line shows the predicted fit.

### Signal strength and responsiveness drive duration

Using the same potency metric, we next tested whether more potent songs evoke longer-duration CSD. Here we were able to incorporate prior playbacks from four additional experiments for a total of 922 postures from 51 birds. We quantified posture duration as the time (to the nearest 1 second) from the initial vertical motion of the longest tail feather to the moment it returned below the level of the body. Across all birds, mean CSD duration was 4.22 (+/-2.58) seconds. Because the mean duration of song stimuli was 1.11s +- 0.113s, behavioral responses often lasted 5x longer than the signal itself. Unlike latency, we did not detect significant individual variation between birds for duration (Figure 3A, ANOVA F=1.686, p=0.19). We used a linear mixed model accounting for bird identity and other batch effects to test for the effect of song potency and block response rate on CSD duration. We found a significant effect between song potency and CSD duration, with higher potency songs stimulating postures of longer duration (Figure 4B, e=1.167, std. error=0.175, p<.0001). Similar to our findings with latency, we also found an effect of female response rate on duration (Figure 3C, e=0.226, std. error=0.1004, p<.05) suggesting that more responsive birds (with a lower response threshold) produce longer-duration CSD.

**Figure 3.**
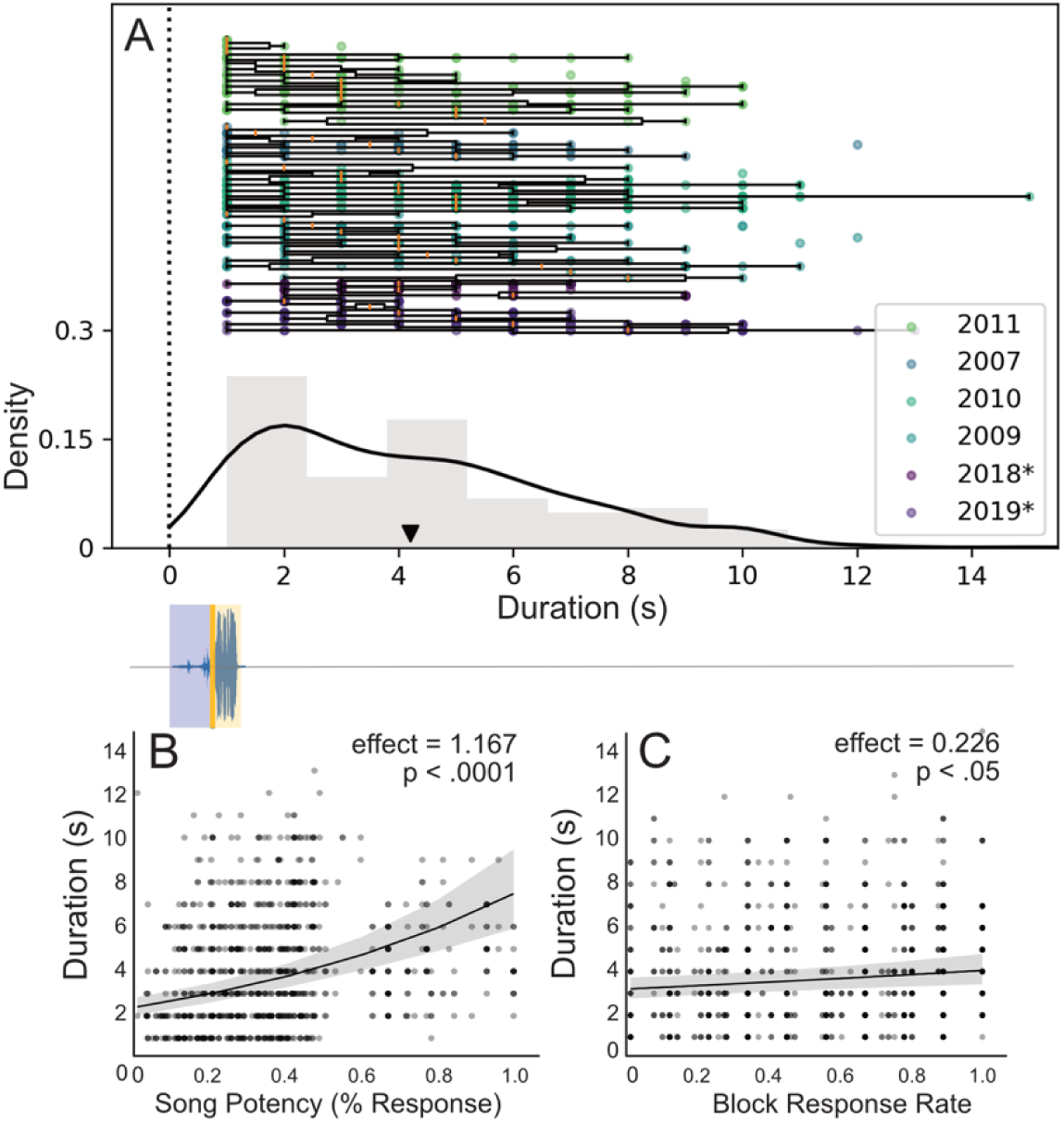
Duration varies with both song potency and underlying receiver state. **A.** Distribution of CSD durations, across 45 birds from 6 experimental groups (n=922 CSDs) aligned to song onset. As in Figure 2, the blue and orange shading show the transition from introductory notes to terminal whistle. Data for 2018 and 2019 are for the same birds shown in Figure 2. **B.** and **C.** show the estimated effect of song potency and block response rate, respectively, using mixed linear models to account for batch effects.

**Figure 4.**
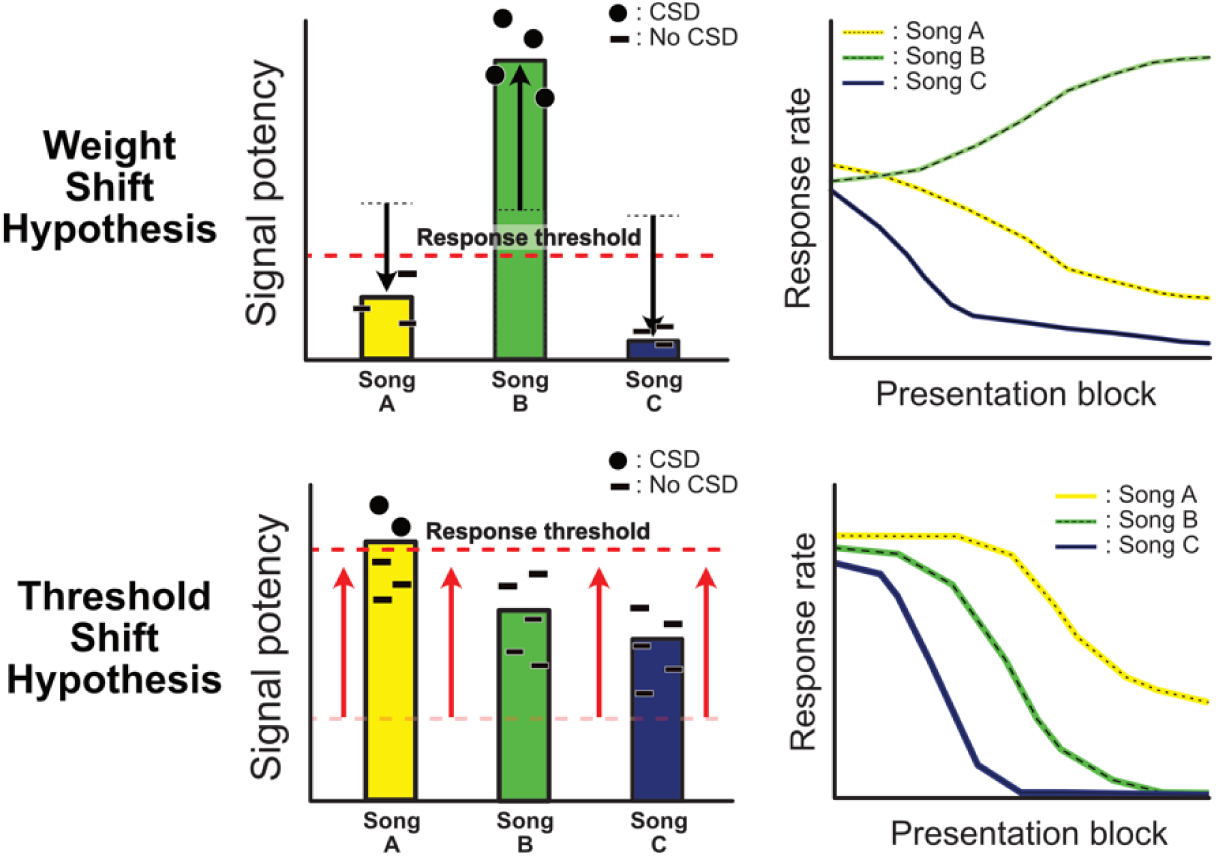
Alternative model mechanisms for the establishment of preferential CSD behavior in a signal-detection paradigm. Both models assume that CSD occurs whenever signal potency exceeds the receiver’s response threshold. The “weight-shift hypothesis” (top left) proposes that responsivity changes to individual songs, via differential weighting of signal strength over time (top right) causing females to increase response likelihood to some songs (e.g. green song) more than others (blue and yellow songs). The “threshold-shift hypothesis” (bottom left) proposes that perceived signal strength for each song is fixed and the response threshold increases over time, such that responsivity decreases for all songs. Females continue to respond (at a lower rate) to the highest quality songs (yellow song) while responses to lower quality songs (e.g. green and blue) disappear entirely.

### CSD trajectory does not vary with responsiveness or song potency

In addition to variation in CSD timing (i.e., latency and duration), we tested whether song potency and receiver state could have meaningful effects on the shape of the pose trajectory itself. Our computer vision allowed us to track the precise trajectory of CSD in 3 dimensions in a select group of birds (n = 208 postures from 6 birds) allowing us to quantify the position and velocity of each body part at each moment in time (Video S2). To define the magnitude of CSD, we used maximum recorded tail velocity and the peak tail height (Figure S2). Unlike timing, we did not observe any differences in the spatial trajectory of CSD posture with no significant effects of either song potency or responsiveness on maximum velocity or peak height (Supplemental Statistics). While these findings do not necessarily rule out posture variation that is too subtle for our approach to capture, we have found no evidence of meaningful variation in the extent of CSD (i.e., tail height), or the velocity of motion by which females arrive there.

It is important to note, that we only quantified pose during “full” CSD, in which the tail was elevated above the body. CSDs (in which the tail never exceeds the level of the body) occurred in response to 9% of songs (compared with 49% for full CSD). Because the tail is partially obscured behind the perch during these partial-CSD, we could not reliably quantify the trajectory of these postures as we did for full CSD, but we could calculate response latency. We observed that partial-CSD latency (mean 1.03s +- 0.29) was significantly slower than full-CSD latency (mean 0.628s +- 0.25), occurring an estimated 252 ms later than full-CSD (linear mixed model, p < .0001), after accounting for individual differences. Unlike full-CSD, we did not find a significant effect of song potency or block response rate on partial-CSD latency, but because partial-CSDs were rare (only 37 examples across 6 birds), our statistical power was low. Interestingly, partial-CSDs occurred most frequently to medium-potency songs (mean potency=0.46, compared to 0.41 and 0.58 for no-CSD and full-CSD playbacks, respectively). As responsiveness decreased over the experiment, reflecting a putative increase in response threshold, the average potency of songs eliciting partial-CSDs increased (effect=0.094, p<0.001;). In other words, less potent songs failed to elicit any response while more potent songs were still able to elicit partial-CSD responses. These partial postures appear to represent “near misses” where the potency did not quite exceed the threshold needed for full CSD.

Taken together, the results in Part 1 suggest that signal potency and female response-rate combine to drive the production of highly stereotyped CSD postures while also influencing latency and duration. Partial CSDs seem to occur when signal potency does not fully exceed female response threshold. This is consistent with a signal-response paradigm where the drive to produce some canonical pose is mediated by receiver state and the strength of the male signal.

### Part 2: Modeling predicts different responses for threshold shift vs song-specific learning

In the experiments described in Part 1, females initially responded to around half of songs (mean = 4.81 out of 10 songs) but appeared to be more selective in later blocks (in the last block only responding to 1.36 songs, on average). This apparent emergence of strong preference for the best songs could be explained by either a “weight-shift” or a “threshold-shift” hypothesis. Here, we develop these hypotheses first as a conceptual model then as a simulation, using a signal detection paradigm, in which behavioral responses occur whenever a stimulus exceeds some internal threshold.

Figure 4 outlines the difference in our two hypotheses: In the “weight-shift hypothesis”, behavioral responses change due to a shifting distribution of (perceived) signal potency. In this scenario, over repeated presentations, the perceived quality of a song changes, relative to other songs, causing a bird to be more (or less) likely to respond. Since in our case, the signal itself is held constant, this would be driven by some form of auditory tuning (Dong and Clayton 2009; Wall and Woolley 2024) by the female, causing the same signal to be perceived differently over time. It is known that this sort of song-specific learning occurs during development and pairbond formation (reviewed in Fujii et al. 2022), and is consistent with how humans learn to recognize and respond to signals in similar experimental paradigms (Salasoo et al. 1985). For this reason we had assumed preference formation observed during CSD was mediated by song specific learning. By contrast, in the “threshold-shift hypothesis”, the relative perceived quality of the signals is constant, but the response threshold itself varies (i.e., birds become more or less responsive in general). Both of these mechanisms can lead to a “best-of-n” ranking and identifying which of these (non-exclusive) mechanisms drives preference formation is an essential aspect of understanding courtship.

### Weight-shift and threshold-shift models can be distinguished by whether song potency diverges

We simulated song playbacks using our two proposed models. In the threshold-shift model, we increase the threshold over the course of playbacks while keeping the random song potency scores constant. In contrast, for the weight-shift model, we keep the threshold constant and vary the song potency values, with all songs starting with equal potency and then gradually shifting (either increasing or decreasing) towards their ranked potency. We used (simulated) experimental blocks (representing one playback of each song within a set) as a representation of time in our model, allowing for comparison with actual playback experiments.

The predictions of the two main models are shown in Figure 5A-B. Both models produce females that initially respond to many songs and eventually limit their responses to a few preferred songs. However, the outputs of these models are visually distinct, and can be differentiated by testing for a divergent song by time interaction (Figure 5A vs 5B). This test successfully distinguishes the two models across all parameters as long as there is least some variation for all songs (Figure S3). In short, the threshold-shift and the weight-shift models can be distinguished based on whether all songs decrease in parallel over time or whether the direction and magnitude of the shift is song-specific, with the most preferred songs eliciting more CSDs—and the least preferred eliciting fewer—over time

**Figure 5.**
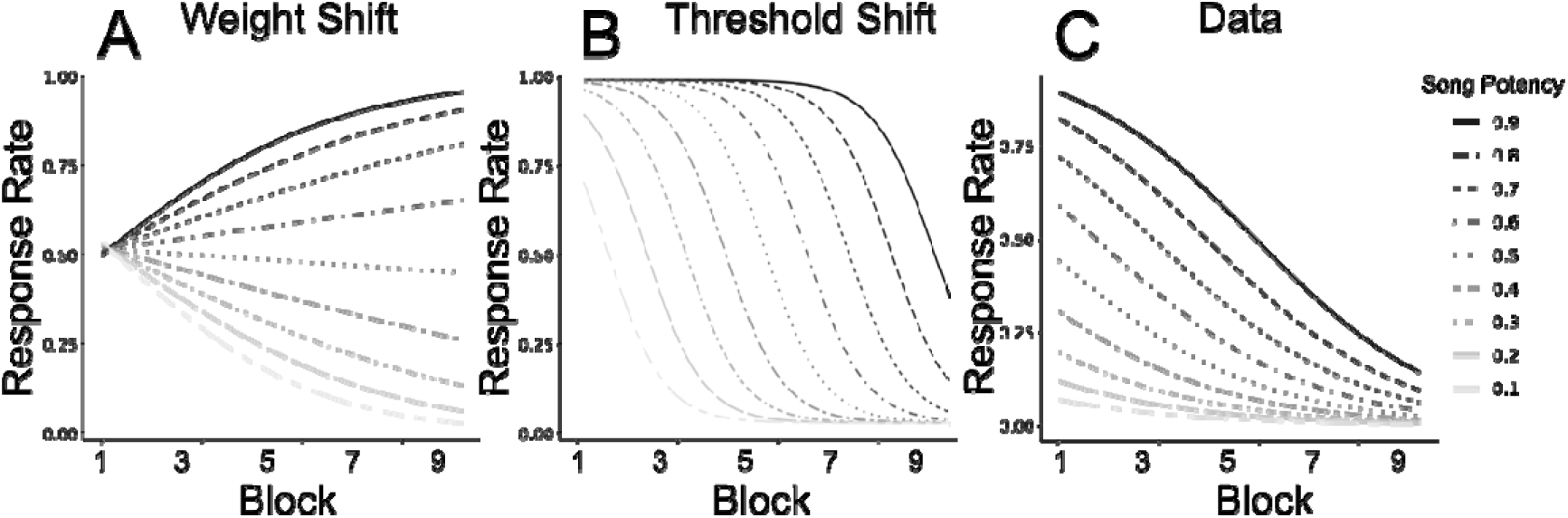
Song selectivity matches the predictions of a threshold-shift model. The left two plots show how response rate is expected to vary over the course of experiments, plotted by song-potency, as predicted by a) a weight shift model, in which individual song-weights vary or b) a threshold-shift model, in which female response threshold varies, or c) as shown in the empirical data. Empirical data are more consistent with a threshold-shift model, in that all songs decrease over the course of playbacks, whereas under a weight-shift model, the highest potency songs are predicted to increase, while the lowest potency song decrease.

While the exact values observed depend on the parameters specified in the model, averaging across all parameters tested, our simulations correctly distinguish the underlying mechanism (weight-shift vs threshold-shift) 98% of the time. In other words, by checking for a divergent song by block interaction, we can distinguish between these two potential mechanisms for a given dataset. It is important to note that these two hypotheses are not exhaustive: signal perception (both in nature and in our model) is subject to noise, and changes in this noise could also shift behavior. We also modeled a “noise-reduction” strategy (Figure S4), in which song weights were constant but noise decreased over time (resulting in perceived potency that becomes more similar to a song’s true potency value). This mechanism also results in a divergent interaction, making it functionally similar to the “weight-shift” hypothesis, such that these two hypotheses would be difficult to distinguish from behavior alone. However testing for divergent song-by-block interaction could rule out both, providing strong evidence for a threshold-shift-like mechanism.

### Shared preferences and rapid responses are consistent with shifting signal response model

Having established these predictions, we applied this test to our experimental data, by aggregating playback experiments recorded from 68 birds, comprising 7 different experimental groups across 10 years. We excluded any bird that produced fewer than 5 CSDs, leaving n=54 birds for our analysis. Comparing our recorded data to the predictions of the model, female responses (Figure 5C) are most similar to the predictions of the threshold shift model (Figure 5A-B). To measure this more precisely, we fit a mixed logarithmic model with bird as a random intercept, and block and song potency as fixed, interacting effects. This model allows for both song potency and time to predict whether a female responds—with the effect of song potency allowed to vary with time—and fits an average responsiveness to each female. Using this model, we found that both song potency (effect: 8.416, p < 0.0001) and block (effect: −0.346, p<0.0001) were strongly predictive of posture, with song potency having a relatively stronger effect within this experiment. Assumed song potency was calculated independently for each bird, based on the responses of other birds (excluding their own responses to prevent statistical confounds). Within our data, high, medium, and low potency songs all decrease steadily over time (Figure 5C) with no divergent song-by-block interaction, consistent with a threshold-shift model (Figure 5B). Additional features of the behavior further support the idea of a shifting threshold: a response in the first block predicted whether a bird would respond in later blocks (effect = 0.175, p < 0.0001), suggesting that relative potency scores were detectable from the first block and remained consistent throughout. Using a generalized linear mixed model including potency and responsivity (as in Part 1), we find that female responsivity, i.e., whether a female responds to the songs immediately before and after a given presentation, is a strong predictor of whether she will respond to a given song (responsivity effect = 5.40, p < 0.0001) although this was not quite as strong a predictor as song potency itself (effect 6.03, p<0.0001). Taken together, this provides strong evidence for a generalized shift in underlying response threshold and rules out a change in the song-specific weighting (at least in this context).

## Discussion

In the current study, we developed a novel approach for quantifying copulation solicitation displays, which were produced in response to varying song stimuli and captured with millisecond precision. We find that female rankings in the CSD assay appear to reflect perceived potency values for the presented songs, the values of which are broadly consistent across individuals and across populations. By quantifying CSD, we show that there is meaningful variation in the timing of CSD, with high potency songs eliciting the shortest onset latencies and longest response durations. Interesting, this response latency was often shorter than the duration of the song itself, meaning females initiated the CSD prior to hearing the terminal whistle (during the note clusters in the first half of the song). Finally, we have shown that the observed change in preferential behavior is more consistent with a gradually increasing threshold (threshold-shift) than learned, comparative rankings (weight-shift). This is further supported by the fact that female preference was consistent across groups, and that CSD usually began during or before the completion of the introductory notes, suggesting an almost-reflexive response to song.

### The all-important response threshold

In our dataset, whether a given signal successfully evokes CSD is a function of both song potency and female responsivity. This implies that potent signals are not a guarantee of copulation, and lower-potency songs can readily accomplish copulation with responsive females. Female response threshold thus appears to act not only as a complete block on copulation at certain times (for example in the non-breeding season) but also provides graded modulation of which signals are sufficient at any given moment during courtship. Threshold-based regulation of selectivity has also been proposed previously in túngara frogs (Akre and Ryan 2011), and may be a consistent mechanism of behavioral modulation, allowing innate responses to be context specific without the need for cognitive control. While some of this attenuation may be explained by short-term sensory habituation (Rankin et al. 2009) it seems unlikely that this fully explains the behavior observed here, since a) song presentations were spaced 90 minutes apart and arranged such that females only rarely heard the same song more than once per day and b) in the wild, females hear multiple songs in quick succession, and courtship often requires long bouts of songs prior to CSD, which is more consistent with a song being stimulating, rather than habituating. Interestingly, auditory responses to song in brain control areas known to influence CSD behavior (Brenowitz 1991; Maguire et al. 2013) have been shown to be strongly gated by the animal’s behavioral state (Schmidt and Konishi 1998), suggesting a possible pathway for linking behavioral responses to the acoustic stimuli. Regardless of the underlying mechanism, the necessity of female responsivity means that courtship is not merely a competition among males to generate the most potent signal possible, but a skillful process where both males and females time their behavior to female receptivity, using a suite of behaviors to identify influence female state.

For the observed link between song potency and both CSD latency and duration means that a male’s window for copulation varies with the potency of song. While this can provide more time for copulation, maximizing CSD duration is not necessarily optimal, as it also opens opportunities for competing extra-pair copulations. This may also explain in part why cowbirds maintain a large repertoire of songs, making it possible for precise control of courtship. Additionally, the importance of receiver state suggests that *when* signals are presented can be just as important as the absolute potency of the signals. The timing of signals is likely to be even more important in natural contexts, where there is presumably more variation in the behavioral state of females.

Given that females, when highly responsive, seem unable to simply “choose” not to produce CSD in response to a potent song, they too likely require strategies to successfully navigate courtship to maintain reproductive control, either by avoiding the song entirely (e.g., by leaving when males approach—the most common response in cowbirds) or by somehow reducing the perceived potency of songs to them. Female cowbirds are known to produce loud, broadband “chatter” vocalizations in response to male song, and these vocalizations have been shown to effectively jam the introductory notes and reduce the chance of CSD, offering a strategy to females for maintaining reproductive autonomy (H. L. Anderson et al. 2021). Even if CSD functions reflexively to male signals, females can use a range of behaviors, including chatter, to avoid, extend, and encourage courtship and copulation. Although the importance of female behavior during courtship generally remains poorly studied compared to male behavior, it is increasingly clear that females act as active participants not just passive receivers (Amy et al. 2015; Akre and Ryan 2011; Rodriguez-Sierra et al. 1975), forcing us to expand our understanding of the roles of males and female roles in courtship.

## Conclusion

We have focused specifically on cowbird copulatory posture, but the mechanisms of copulatory control are likely shared—not just across songbirds, but possibly throughout birds and other animals producing similar lordosis responses (Perkes et al. 2019). Our approach here provides a template for exploring postural behavior that can be applied in future studies. As we have shown, CSD can be studied in controlled, quantifiable, high-throughput approaches. It thus serves as an excellent model of copulation specifically and postural control generally. By pairing theories of action regulation with detailed postural quantification as demonstrated here, we can begin to identify the critical processes the regulate copulation and successful reproduction.

## Materials and Methods

### Subjects & Housing

A total of 83 female cowbirds were used in these experiments—including the data gathered from previous experiments. Some birds did not meet the inclusion criteria for specific approaches and were thus only included for a subset of analyses (Table S1). All birds were captured in Pennsylvania (in Chester or Montgomery Counties) between the years of 2007 and 2018 under appropriate state and federal trapping permits, using funnel traps. To acclimate to captivity, females were captured at least 6 months prior to playback experiments (during the fall of the prior year) and then cohoused in pairs or small groups. During this habituation, and when after participating in playback experiments, birds were housed either in small flocks in aviaries (at least 1.2×1.2×2.4m)) or co-housed in smaller wire cages (56cmx56cmx46cm). All housing was supplied with adequate perches and birds had access to food (a modified Bronx Zoo diet for omnivorous birds mixed with seed) and water, *ad libitum*.

At least one month prior to the start of playbacks, males were removed from the room/aviary in which females were housed in order to prevent female exposure to male song and promote CSD responses to playback (King and West 1983). Based on the capture date and time spend in captivity, females were at least 1 year old during playbacks.

### Acoustic Stimuli

All songs used as stimuli were gathered from captive males in aviaries, using a boom mic. Cowbirds have a distinct “perched song”, lasting roughly 1 second, consisting of 2-3 low, burbly note clusters, followed by a high-pitched whistle (Figure 1, Supplemental Video S1). Both the note clusters and the whistle have been implicated in song potency. Although male cowbirds also produce other vocalizations, these perched songs are by far the most common signal directed to females, and all acoustic stimuli used for playbacks were of this form. In all but two groups, we used a single set of 10 songs recorded from aviaries in Montgomery County, PA in 2008 (Maguire et al. 2013). In 2007 we used a different set of songs, also recorded in aviaries from local males (Ronald et al. 2017). In 2009 we used songs recorded under controlled conditions as part of a separate experiment (Gersick and White 2018). Acoustic quality across songs was similar, and songs from different groups were never mixed. All stimuli were most likely new to the experimental females, since we used females who had not been housed with the males that produced the songs, though we cannot rule out the possibility that females caught between 2007-2009 had interacted with the males prior to capture.

### Playback Procedures

We follow a previously described procedure for CSD assays using song playbacks to female cowbirds (King and West 1983; White 2010). Females, overwintered in captivity, and isolated from males during breeding season often (but not always) produce CSD in response to song playback, allowing the number of CSDs produced to be used as a metric of “potency” for each song (Figure 1), representing the average probability that a given will evoke a CSD in this context. Prior work has shown that the measured song potency accurately predicts courtship success (O’Loghlen and Rothstein 1995; West et al. 1981). In our study specifically, females heard the playback of a single song, played over a speaker. Speaker parameters and distance varied slightly between (but not within) experimental groups, but sound intensity was maintained between 80 and 90 dB (measured using a sound level meter placed at the location of the perch.) When possible, we waited to play song until females were stable on the perch, however if this did not occur within the 5 min playback window, we played the song regardless. We presented one song every 90 minutes (to minimize habituation), for a total of 6-8 songs per day. Songs were arranged in pseudorandom blocks, such that females heard every song in the set before repeating. Playback experiments lasted 2-4 weeks per bird (depending on the number of songs and repetition blocks, see Table S1). We did not use any hormonal manipulations (e.g., estradiol implants). Our dataset can be divided into three experimental groups, for which we detail the specific housing and playback procedures below (see also Table S1).

### Multi-camera cages

During our experiments performed in 2018-2019, we used a custom cage (55×55×48cm) equipped with an array of 4 cameras allowing us to define CSD pose with high temporal and spatial precision for n=15 birds. This cage was contained within a sound isolation chamber (76×76×76 cm, adapted from 749 instructions from accousticsolutions.com). Within these chambers, we performed playback experiments following the general setup described above. In this case, recording and playback were automated using a custom python script. We estimated the position of the bird in real time using a color discriminator to detect the bird, so that we could wait to play the song until the bird was resting on the perch for 10s (if this condition was not detected within 5 minutes, a song was played regardless). We used four FLIR Grasshopper color cameras which could be synchronized via a hardware trigger. We were able to record from all four cameras, 50fps at 1080p, providing high temporal and spatial resolution. We performed intrinsic calibration (measuring the sensor and lens properties of the camera) once with a printed checkerboard prior to the experiment (https://github.com/ros-perception/image_pipeline.git). For extrinsic calibration (identifying the relative position and orientation of the cameras), we carefully measured the positions of each camera, and also placed AprilTag markers (Olson 2011) on the four walls of the cage, allowing us to account for small variations due to mechanical perturbations or thermal expansion using TagSLAM software (Pfrommer and Daniilidis 2019).

We quantified bird pose using the automated tracking program SLEAP (Pereira et al. 2019), employing a single-instance model (based on a pretrained efficientnetb3 model (Tan and Le 2019). This provided 2d keypoints in each camera, which we triangulated based on the calibration described above. Since most of the motion CSD occurs in the vertical elevation of the tail and body, we simplified the representation during CSD as the vertical distance from the tip of the tail to the feet (Figure S2). To reduce the effect of noise from mis-identified frames, after 3D positions, keypoint trajectories were smoothed using a 1d gaussian kernel with sigma=5. Based on this trajectory, we also calculated the <u>maximum tail velocity</u> (in the 2s window following onset), and the peak tail height, in order to define the extent of the posture.

### Cage rack playbacks

In 2019, to increase the throughput of playbacks, we developed a method to play songs to females housed in standard wire cage racks, rather than in individual enclosures. During May and June, we housed 18 adult females in 9-cage wire racks [a 3×3 rack of cages, each measuring 56 x 56 x 46 cm, with one female per cage]. We performed these experiments for two consecutive recording blocks. Videos were recorded using four Logitech C920 HD webcams (the multiple cameras were necessary to cover all 9 cages). A custom python script automated the playback and video capture. The script tracked bird motion and attempted to wait until total motion was low, but would play a song regardless after 1 minute.

This allowed us to play song to multiple females simultaneously to generate a larger dataset of CSD responses, but without the temporal and spatial resolution of the multi-camera recordings, meaning it was impossible to precisely define CSD latency or duration.

### Cohoused playbacks

To further increase our sample size, we used existing data from an additional 50 birds from previous playback experiments. These playbacks were conducted between 2007-2011. In two of these experiments, we used a set of songs that was different from the 2008 song list used in all other experiments (Table S1), but the design of these experiments was otherwise the same. In all cases, females were housed in pairs within sound attenuation chambers. Cohousing has been shown to reduce stress and has been shown to have no influence on playback responses (Smith et al. 2000; Eastzer et al. 1985; King and West 1983). Every 90 minutes, human observers confirmed that a female was on their perch before playing the song stimulus. If the female never settled on the perch, the song was played regardless.

This allowed for quantification of CSD production and duration (to the nearest 1 second), but we did not have the precise video-acoustic calibration necessary to easily quantify latency.

### Behavioral scoring

In all experiments, video recordings were scored by trained observers. We noted CSD (including full or partial, described below) and recorded duration, except in the case of the cage-rack playbacks where video resolution and frame rate precluded measuring duration with confidence. During the posture-analysis experiments, human observers also manually identified female responses to male song playbacks. We defined **CSD** (also called “Full-CSD”) as arching of the back to elevate the head and tail feathers while spreading the wings. To qualify as CSD, tail feathers needed to be raised above the body. A display was defined as *“***partial-CSD***”* if the female initiated the characteristic wing spread and/or back arching, but did not elevate her tail feathers above the body. CSD duration was calculated as the time (in seconds) between initiation and the moment tail feathers descended below the body after reaching the peak. Latency was defined as the time between the start of the song playback, and the first upward motion of the tail feathers.

### Potency and Responsivity

Song potency is defined as the average rate at which all females respond to a given song, while responsivity is the average response rate of a given female to all songs, calculated on each day of the experiment. In order to avoid statistical confounds, potency is calculated on a per-bird basis, using all other birds (excluding the focal female) to calculate potency. Similarly, responsivity at the moment of the playback is calculated as the proportion of songs to which a female responds on a given day, excluding the playback itself.

### Data inclusion and statistical analysis

Analyses can be divided into 3, roughly nested groups of birds: latency/pose, duration, and CSD production (Table S1). For latency and pose (i.e., max tail height, tail velocity), we included birds from the multi-camera cage who produced CSD while on a perch (n=8 birds), with n=6 birds producing partial CSD on the perch. For duration, we excluded birds in the cage rack playbacks (for whom duration could not be determined) and any other birds who did not produce CSD, leaving n=51 birds. For assessing CSD production, we included any bird that produced > 5 CSDs, for n=53 birds. For analyses, we primarily used linear mixed models to account for individual, block, and group-level effects. The specifics of these models, along with other statistical tests used, are described in the relevant results sections, with full details in the supplemental statistics section.

### Permutation analysis for preference

To test whether birds showed a significant preference for songs, we performed a permutation test where we shuffled within-bird responses, then calculated averaged (shuffled) responses for each song, sorted them, and calculated the slope. We repeated this procedure 10,000 times to generate a distribution of expected sorted slopes if song responses were random. For assessing the strength of various effects on CSD production, we used linear and generalized mixed models with song potency and block as fixed effects, with bird ID, song, and group as random intercepts. As the independent and response variables used are specific to each section in the results, the details of these models are described as part of the relevant results section and Supplemental Statistics (which contains the full output). For behavioral quantification and data processing, we used python 3.7. Statistical tests were performed using R (version 3.4), either directly or via the python wrapper Pymer4.0 (Jolly 2018)). Plots were generated in python using matplotlib or in R using ggplot. Regression plots show the estimated effect and 95% confidence intervals. Descriptive statistics are written as mean +/-standard deviation unless otherwise noted.

### Signal-detection model and simulation

To generate specific predictions for testing comparing the importance of song potency and female responsiveness, we developed a simulation using a signal-detection model, varying either the response threshold or signal weights over the course of the experiment comprising 10 birds, exposed to 10 songs, over the course of 10 blocks, repeated for 100 iterations. This represents a manageable number of birds and playbacks for a given experiment, since real experiments are limited by the space and time available to perform playbacks. All threshold and potency values were between [0,1]. For each presentation, a potency value was generated with some random uniform noise ([-0.1,0.1]) around the intrinsic potency value of that song in the block. Song potency values could be random or highly stratified, based on the “bias” parameter. For the signal-weight model, threshold was fixed (between 0.25 and 0.75) while initial potency was 0.5 and then transitioned evenly to the ranked-potency values ([0.05-0.95]) according to “learning-rate” (set between 1 and 10 blocks). For the threshold-shift model, song-potency scores were held constant at their ranked-potency values, while the threshold transitioned evenly from 0.25 to 0.75. At the time of a presentation, if the song-potency value with some “error”, set between [0.0,0.5], exceeded the current threshold, a CSD occurred. Thus, we had four variable parameters (learning rate, threshold, error, and bias) over which we iterated. The model was implemented in python (3.7), which output simulated datasets which we analyzed using the same approach as for assessing preference in our real dataset, described above.

### Data Availability

All original CSD videos, along with all meta information, human annotations, and reconstructed postural trajectories, along with the processing and analysis code, are archived on Zenodo (10.5281/zenodo.22101255).

## Supporting information

Supplemental Statistics

## Acknowledgments

We wish to acknowledge Nikos Kolotouros for his help with the triangulation code, and Jakub Jarmula for his tireless assistance building cowbird housing. 3D-printed perches were courtesy of the University of Pennsylvania Libraries’ Biomedical Library. This work was supported by an NSF PRFB 2209270, and IOS:1557499

## Supplemental Figures and Table

**Figure S1.**
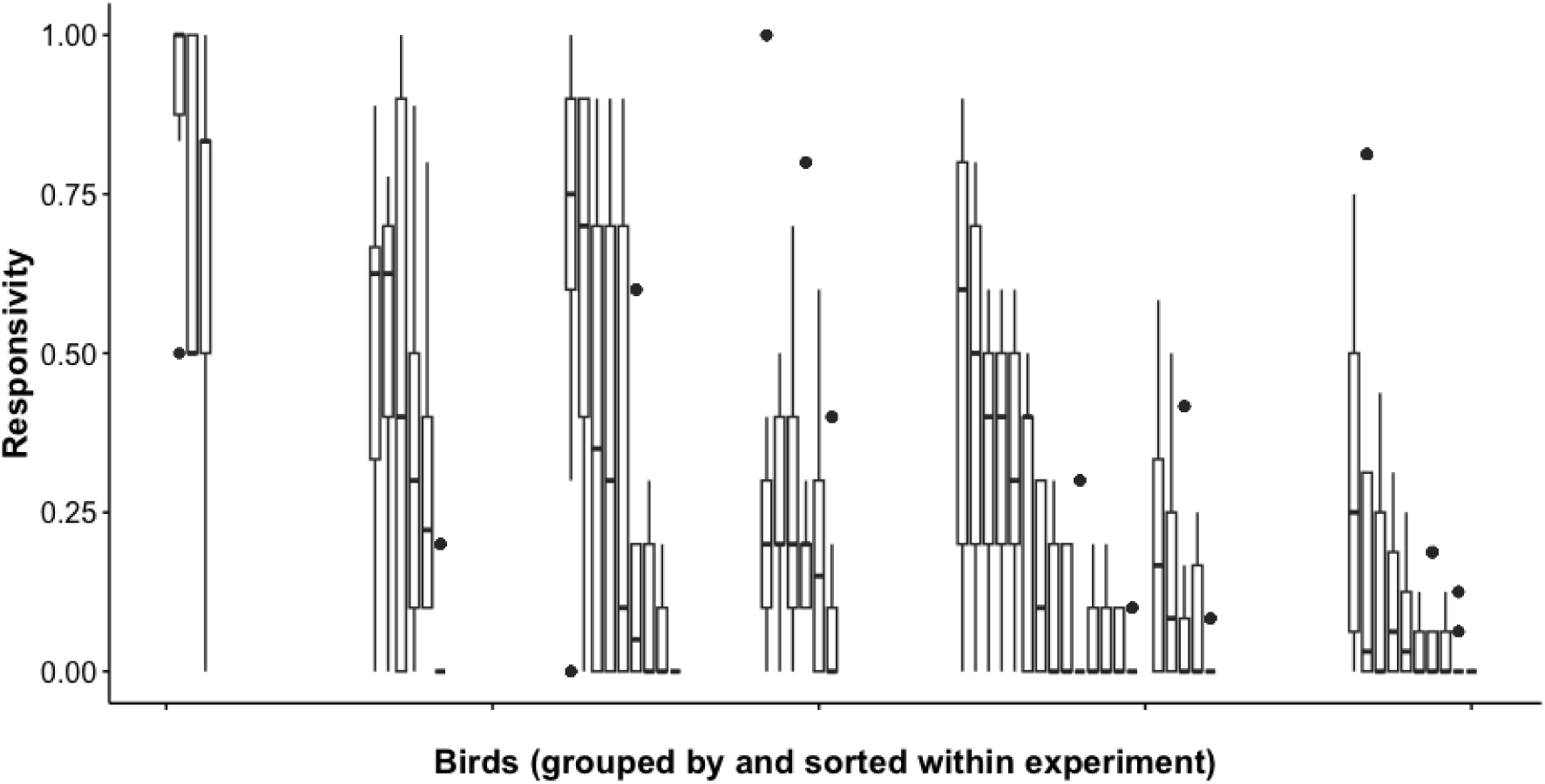
Individual variation in responsivity. Each box plot shows the distribution of mean block response rate for an individual bird. Birds are divided into experimental groups, and sorted within-group by median response rate.

**Figure S2.**
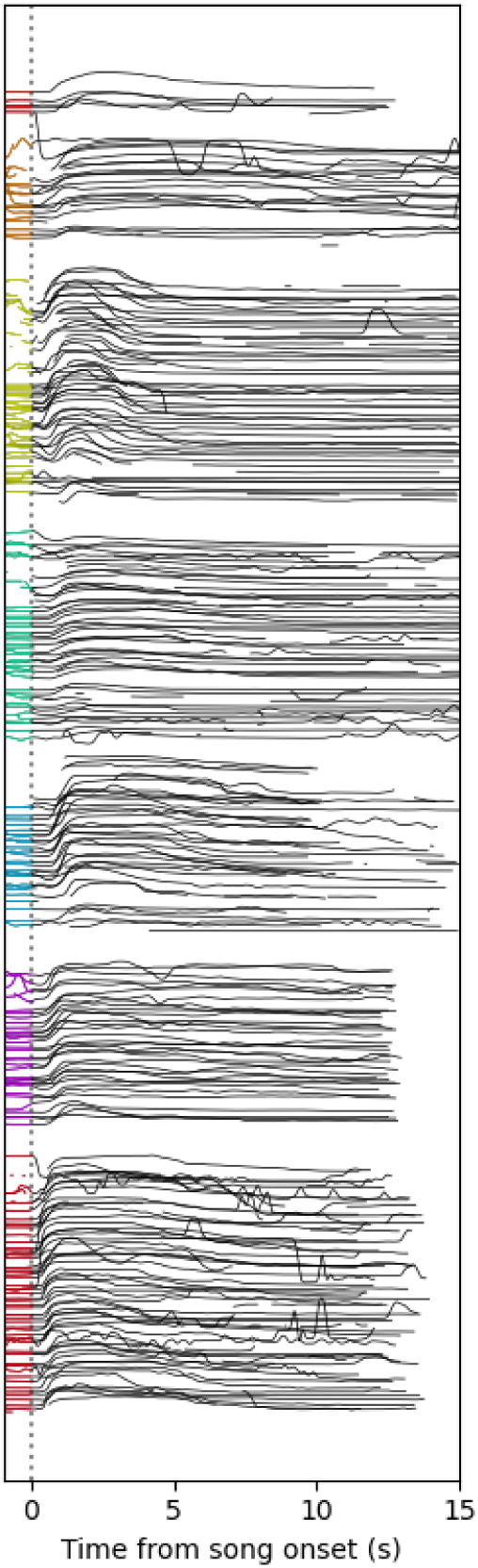
Overlayed plots of all CSD trajectories. Each line shows the trajectory of an individual CSD, plotted as tail height (normalized to the position of the feet). Trajectories are divided by bird (shown by the different colours of the line prior to song onset). Within-bird, trajectories are sorted by latency, from shortest to longest.

**Figure S3.**
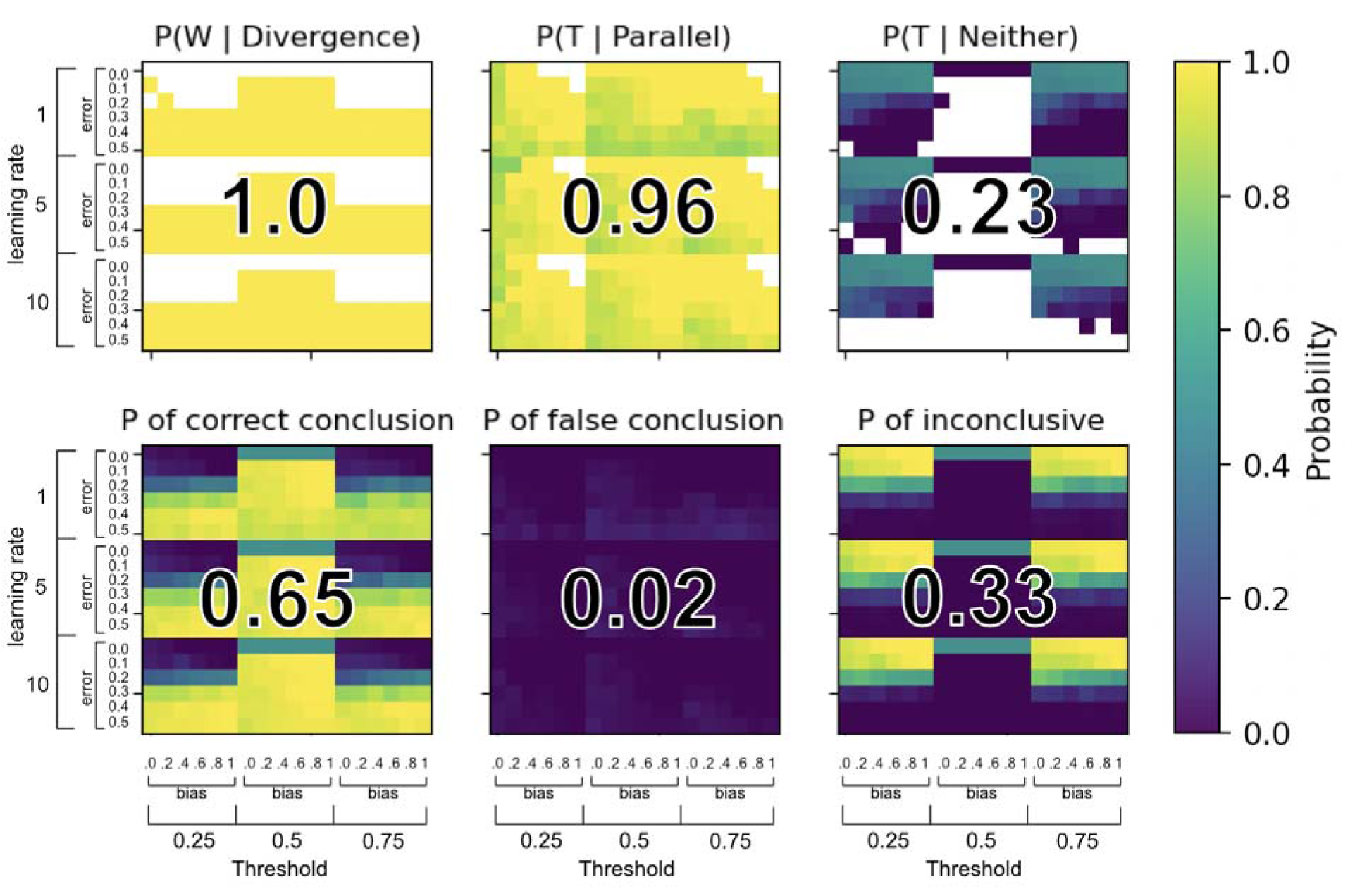
Model Parameter Exploration. These heatmaps show the parameters under-which we can correctly infer the underlying model, given that the simulated data shows divergent vs parallel slopes for the most- and least-potent songs. Overlayed numbers show the mean probability for a) the probability of a weight-shift model, given divergent response rates; b) the probability of a threshold-shift model, given a parallel response rates; c) the probability of a threshold-shift model, given a null result (usually meaning the top or bottom songs do not vary over the course of the experiment). Using bayes theorem, assuming either model is equally likely, we can calculate d) the probability of obtaining correct conclusion, e) the probability of obtaining a false conclusion, or f) the probability of obtaining an inconclusive result. Axes show the four parameters over which we iterate, defined in the methods section.

**Figure S4.**
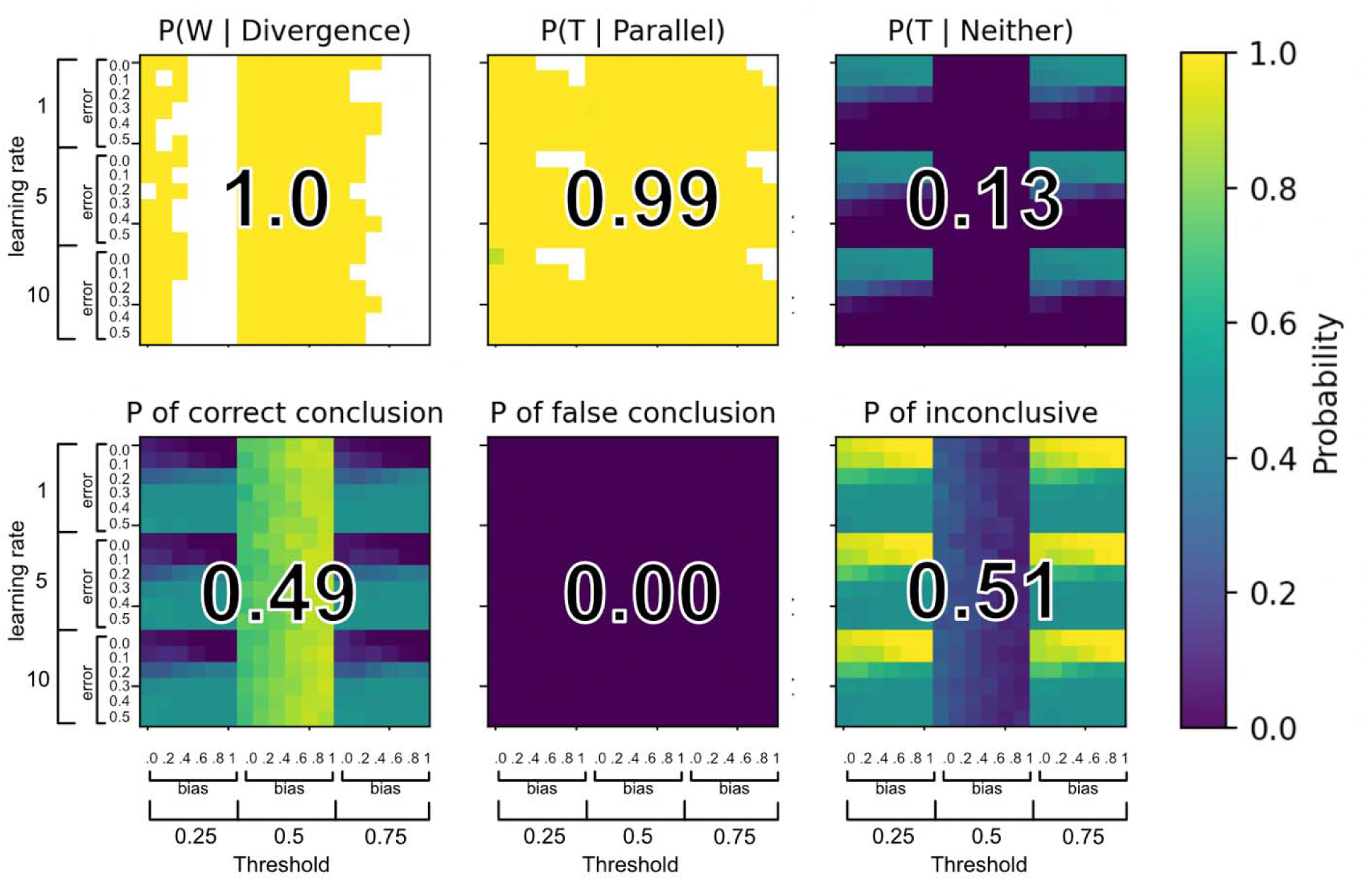
Noise-shift Parameter Exploration. This plot is identical to Figure S3, except instead of a weight-shift model, the threshold-shift model is compared against a noise-shift model, in which the variance of the song-quality distributions decreases over time, suggesting that the bird’s perception of the song more closely represents its “true” potency. In the noise-shift model, learning rate is the number of rounds it takes to arrive at 0, while error has no effect on noise-shift (since it always starts at 0.5). Potency values do not vary within a given simulation for either model.

**Table S1.** Summary of birds used for experiments. The values denote the specific number of birds used for a given analysis from a group of birds.

| Group ID | Source | Year | n Birds | n Birds analyzed | Pose | Latency | Duration | Responsivity | Song dataset | Notes |
| --- | --- | --- | --- | --- | --- | --- | --- | --- | --- | --- |
| 0 | Multicamera | 2018 | 8 | 3 | 3 | 3 | 3 | 3 | 2008 |  |
| 1 | Multicamera | 2019 | 7 | 7 | 3 | 5 | 6 | 6 | 2008 |  |
| 2 | Cagerack | 2019 | 18 | 8 | - | - | - | 6 | 2008 |  |
| 3 | Cohoused | 2009 | 11 | 11 | - | - | 6 | 5 | 2009 | Previously Gathered |
| 4 | Cohoused | 2010 | 14 | 14 | - | - | 12 | 10 | 2008 | Previously Gathered |
| 5 | Cohoused | 2007 | 10 | 10 | - | - | 9 | 9 | 2007 | Previously Gathered |
| 6 | Cohoused | 2011 | 15 | 15 | - | - | 15 | 14 | 2008 | Previously Gathered |
| X | Multicamera | 2017 | 4 | - | - | - | - | - | 2008 | Pilot Birds |
| Total | 3 | 7 | 87 | 68 | 6 | 8 | 51 | 53 | 3 |  |

