## Supplemental Statistics for "Shifting sensitivity and signal-dependent timing in the copulatory displays of songbirds"

CSD Supplemental Statistics.

Full model outputs for stats found in the main body. See also R.script

Latency

Latency ANOVA

Model Diagnostics

```
> summary(resLat.aov)
             Df Sum Sq Mean Sq F value Pr(>F)
Bird          1  4.115   4.115   88.34 <2e-16 ***
Residuals   266 12.392   0.047
---
Signif. codes:  0 ‘***’ 0.001 ‘**’ 0.01 ‘*’ 0.05 ‘.’ 0.1 ‘ ’ 1
```

Latency Regression

```
Linear mixed model fit by REML. t-tests use Satterthwaite's method ['lmerModLmerTest']
Formula: Latency ~ AvgPotency + BlockResponseRate + (1 | Bird) + (1 |
    Song) + (1 | Block)
   Data: lat_data

REML criterion at convergence: -181.5

Scaled residuals:
    Min      1Q  Median      3Q     Max
-3.3978 -0.6039 -0.1083  0.5214  3.2721

Random effects:
 Groups   Name        Variance  Std.Dev.
 Block    (Intercept) 0.0008939 0.02990
 Song     (Intercept) 0.0011894 0.03449
 Bird     (Intercept) 0.1004904 0.31700
 Residual             0.0240207 0.15499
Number of obs: 268, groups:  Block, 20; Song, 12; Bird, 8

Fixed effects:
                   Estimate Std. Error        df t value Pr(>|t|)
(Intercept)         1.02160    0.12470   9.11335   8.193  1.7e-05 ***
AvgPotency         -0.22994    0.07057  20.07507  -3.258  0.00392 **
BlockResponseRate  -0.11254    0.05031 185.88898  -2.237  0.02649 *
---
Signif. codes:  0 ‘***’ 0.001 ‘**’ 0.01 ‘*’ 0.05 ‘.’ 0.1 ‘ ’ 1

Correlation of Fixed Effects:
            (Intr) AvgPtn
AvgPotency  -0.328
BlckRspnsRt -0.249  0.067
```

Model Diagnostics

We use the package DHARMa (citation), in addition to traditional regression and qq-plots, to assess model fit. In this case, while QQ and residual plots look reasonable, DHARMa shows that the model is underdispersed (dispersion = 0.50603, p-value = 0.232), which likely means the model is too complex and somewhat overfit, weakening our statistical power (This is somewhat unexpected, since BlockResponseRate and AvgPotency already account for our random effects.)

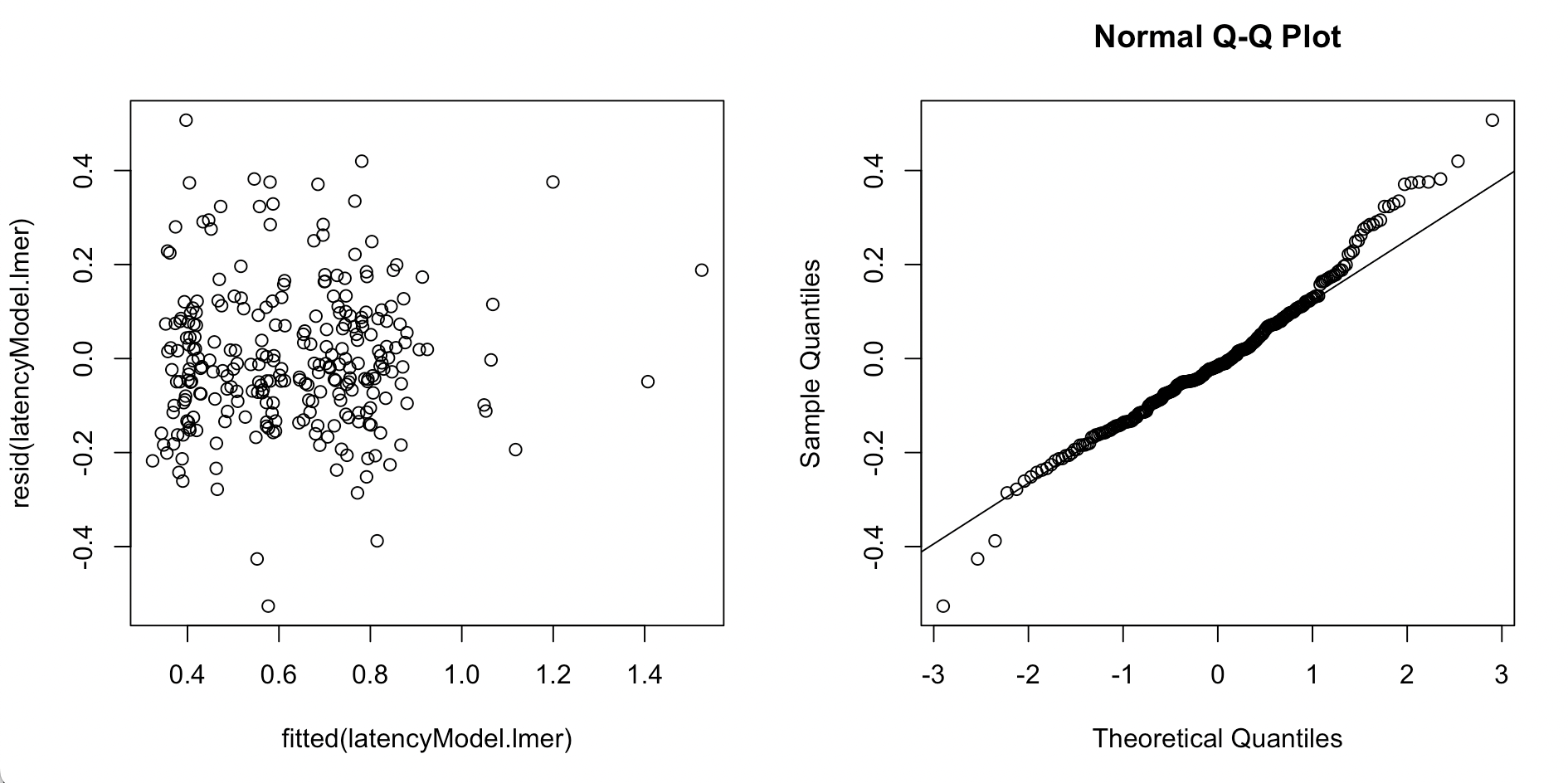

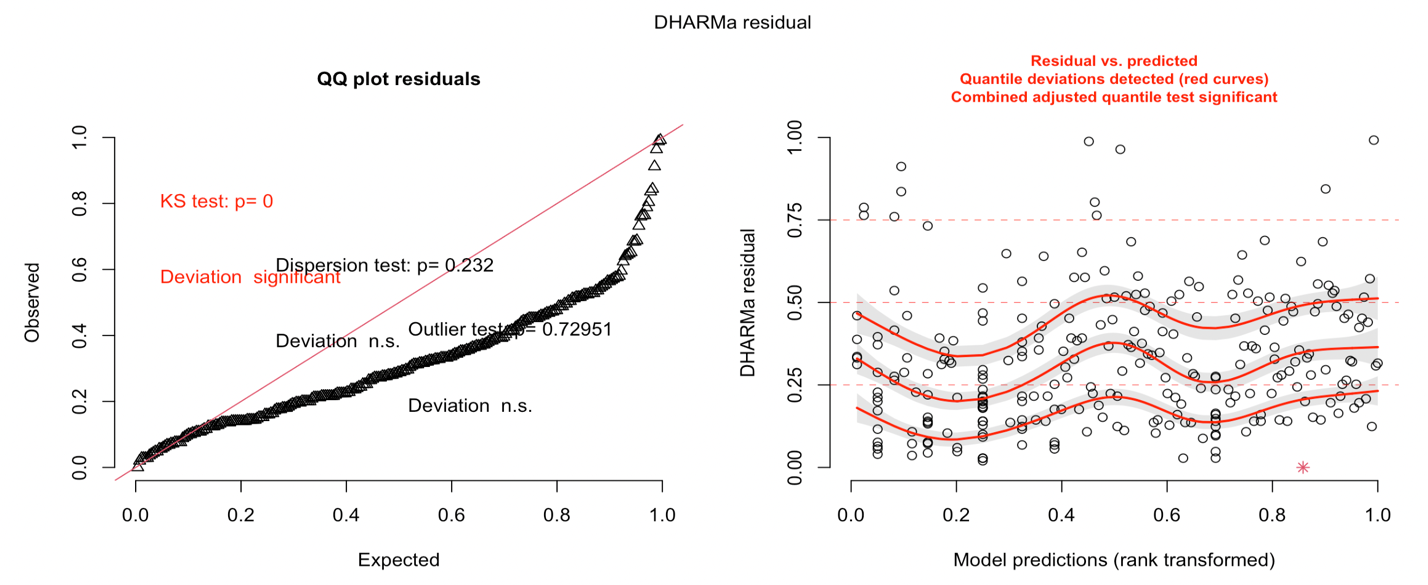

```
Since our results are significant, low power is not concerning, however we also tried a simpler model, which was evenly dispersed, and for which the results were similar (see below).

Simpler model:

Call:
lm(formula = Latency ~ AvgPotency + BlockResponseRate, data = lat_data)

Residuals:
     Min       1Q   Median       3Q      Max
-0.76444 -0.14136 -0.00999  0.13544  0.81766

Coefficients:
                  Estimate Std. Error t value Pr(>|t|)
(Intercept)        0.97986    0.04123  23.763  < 2e-16 ***
AvgPotency        -0.16010    0.05550  -2.885  0.00424 **
BlockResponseRate -0.38150    0.04616  -8.265 6.76e-15 ***
---
Signif. codes:  0 ‘***’ 0.001 ‘**’ 0.01 ‘*’ 0.05 ‘.’ 0.1 ‘ ’ 1

Residual standard error: 0.2148 on 265 degrees of freedom
Multiple R-squared:  0.2591,	Adjusted R-squared:  0.2535
F-statistic: 46.34 on 2 and 265 DF,  p-value: < 2.2e-16
```

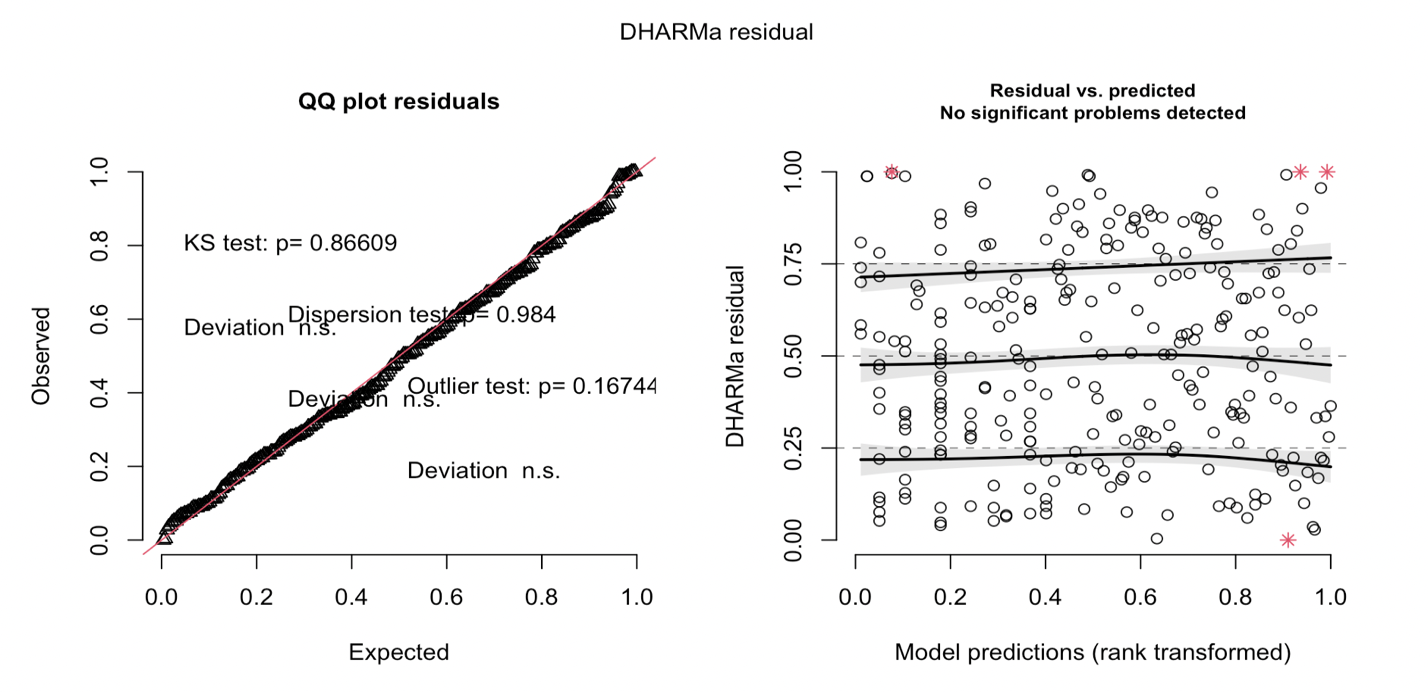

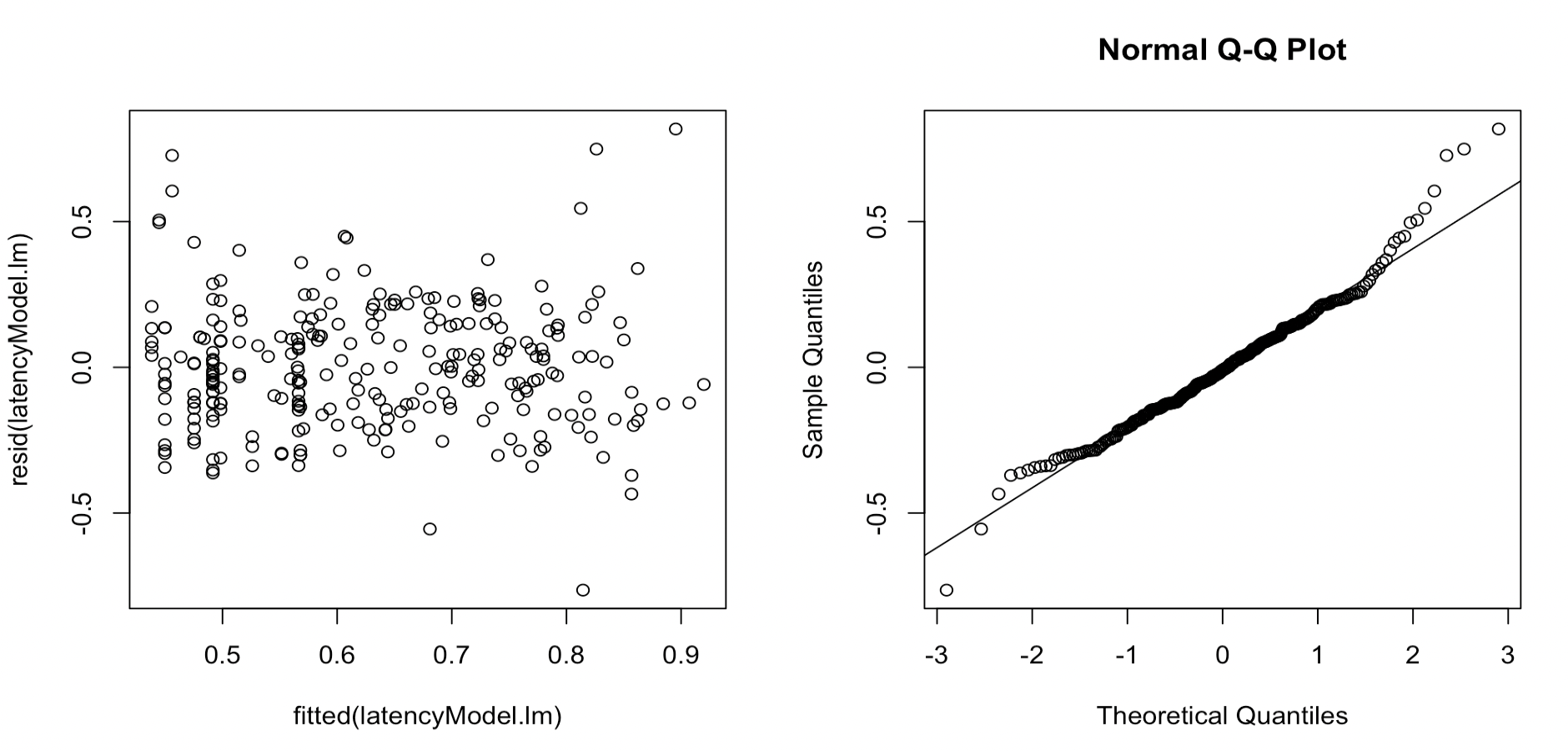
